# Evaluating home-cage-activity metrics for detecting postprocedural pain in a plantar incision mouse model

**DOI:** 10.64898/2026.09.23.753854

**Authors:** Morten E. Malmberg, Tobias Bertelsen, Peter Bollen, Klas Abelson, Sara Hestehave, Otto Kalliokoski

## Abstract

Detection of postoperative pain in mice is challenging, and there is a need for unbiased, automated methods. Home-cage monitoring systems are promising, but their ability to detect mild pain remains unclear. We evaluated whether home-cage activity measures can be used to detect pain-related changes in a plantar incision model of postoperative pain. First, we confirmed the presence of local hypersensitivity using von Frey filaments on 48 C57BL/6JRj mice (24 males, 24 females). Subsequently, we pair-housed 64 mice (32 males, 32 females) and allocated them to receive either a 4 mm plantar incision on the hind paw under anaesthesia, or anaesthesia alone. Baseline locomotion and voluntary wheel-running activity were recorded over five days, followed by four days of postprocedural monitoring in the home cages. Both procedure groups (incision and anaesthesia-only) showed reduced locomotion and running wheel activity relative to baseline. However, activity measures did not differ between incised and anaesthesia-only mice, and we detected no sex-related effects. Under these conditions, home-cage activity measures did not reliably discriminate pain-related behavioural changes in the model, despite the model showing local mechanical hypersensitivity in the von Frey test. Together, these findings indicate that, given the mild and localised nature of the plantar incision mouse model, commonly used home-cage metrics cannot yet reliably detect subtle pain-related behavioural changes.

## Introduction

Detecting postoperative pain in laboratory mice is challenging^1^. Anaesthesia and surgical procedures can alter normal behaviour, but mice may actively mask signs of discomfort, thereby obscuring behavioural indicators of pain^2, 3^. The von Frey test is commonly used to assess pain-related changes by measuring mechanical sensitivity. By applying calibrated monofilaments to an area of interest, the presence or absence of a withdrawal responses can be used to assess analgesic effects or allodynia. However, several factors may influence the assay outcome, including the sex of the experimenter^4^, the surface on which the animal is tested^5^, home-cage bedding material^6^, and whether the animals are stressed^7^. The assay also requires prolonged acclimatisation prior to each test, is prone to observer bias^8^, and necessitates repeated handling to obtain time-course data, which may further confound the measurements. Finally, because the von Frey test captures only a brief snapshot of local mechanical sensitivity, it does not reflect the animal’s overall state before, during, nor after the test.

Recent technological advancements enable continuous assessment of animals in their home cages around-the-clock. Quantification of non-evoked, spontaneous behaviour under undisturbed conditions in the home cage provides observer-independent insights into activity patterns that are not readily detected by brief, experimenter-evoked tests^9^. A key strength of these home-cage monitoring (HCM) systems is their capacity to continuously collect behavioural data without human intervention. These systems could therefore contribute to refinement of welfare monitoring, for example by facilitating earlier detection of adverse reactions to experimental conditions, including surgery-related effects. However, the sensitivity of current HCM-derived metrics to detect mild postoperative pain-related behavioural changes has not been established.

In this study, we examined whether home-cage monitoring detects pain-related behavioural changes in mice following plantar incision surgery, a model of mild postoperative pain.

Given the continued overrepresentation of male animals in pain research^10^, and evidence for sex-dependent responses to noxious stimuli^11^, both male and female mice were included.

We tested four hypotheses: We expected HCM-derived behavioural measures to detect pain-related changes after surgery. These changes were predicted to be most pronounced during the immediate postprocedural period, defined as the first 24 hours following recovery from anaesthesia. We further hypothesised that HCM-derived pain-related behavioural changes would not differ between male and female mice. Finally, we expected running-wheel activity to be more sensitive than cage-level locomotion activity for detecting pain-related changes in the plantar incision model.

## Results

### Evoked behavioural responses

To validate the plantar incision model and confirm measurable evoked behavioural changes in our experimental setup, von Frey monofilament testing was performed (Fig. 2). Mice were assessed once daily for five days following isoflurane anaesthesia with or without plantar incision (Fig. 1a). Analysis of variance (ANOVA), excluding preprocedural baselines, with time as the repeated factor, procedure (incision/control) and sex (male/female) as between-subject factors revealed increased postprocedural mechanical sensitivity in incision animals (F(1, 44) = 42.8, P < 0.001). Neither time, sex, nor any interaction effects were significant.

**Fig. 1:**
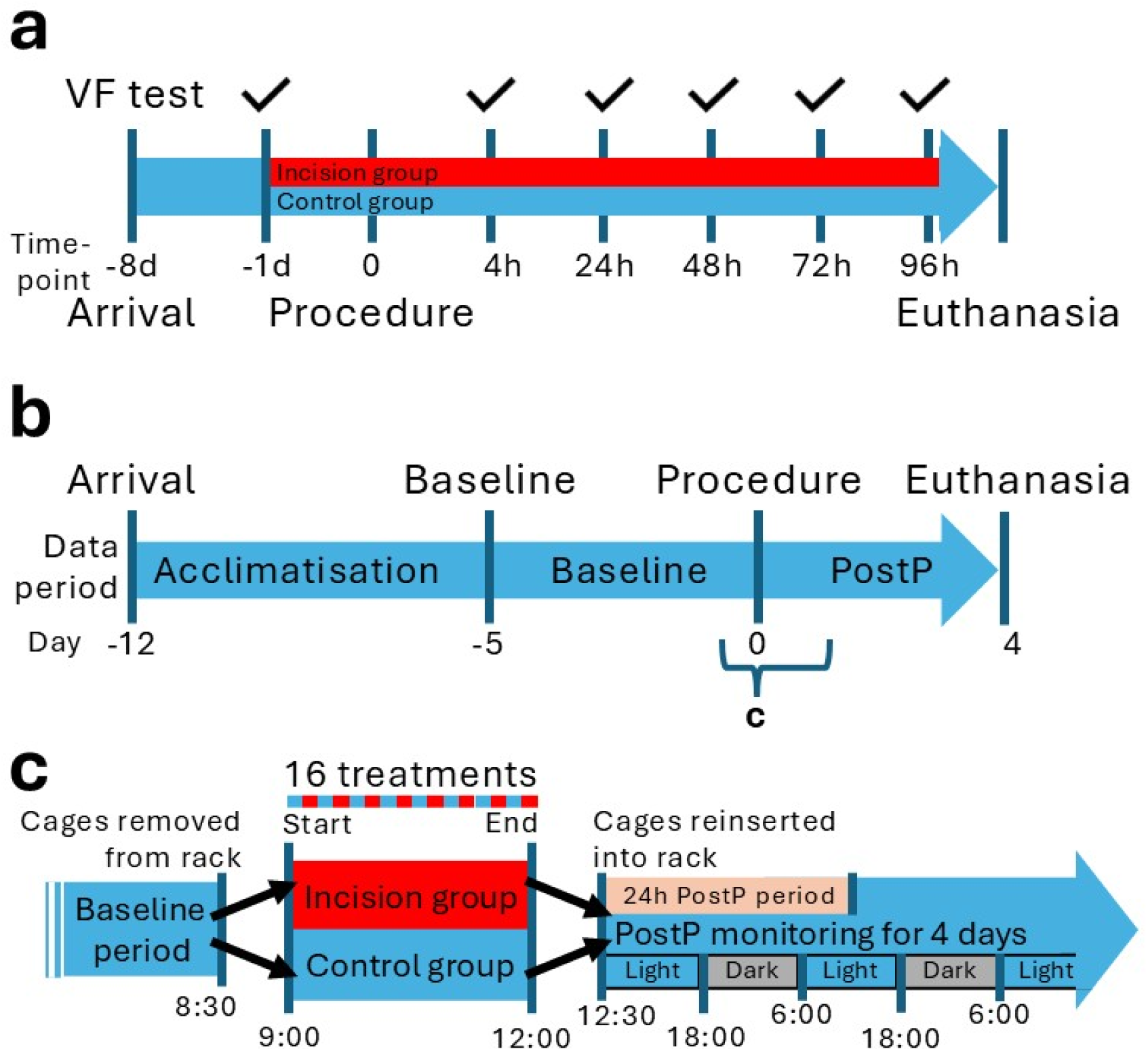
Experimental design and timeline of the two experiments. **a**, Timeline of von Frey (VF) study, with VF tests at the timepoints marked with a tick, where d represents day prior to the procedure and h represents hours after the procedure. **b**, Timeline per cohort of the home-cage monitoring experiment with periprocedural period further elaborated in panel c. PostP = postprocedural monitoring period. **c**, Periprocedural timeline and experimental design. The baseline period ends when cages are removed from the monitoring rack prior to the procedure. After mice were treated sequentially with anaesthesia and cleaning of the right hind paw (control), and the incision group also received an incision, all cages were simultaneously returned to the monitoring rack, marking the start of the postprocedural (PostP) monitoring period. The immediate 24-hour postprocedural period starts data collection at ∼12:30 and ends 24 hours after.

**Fig. 2:**
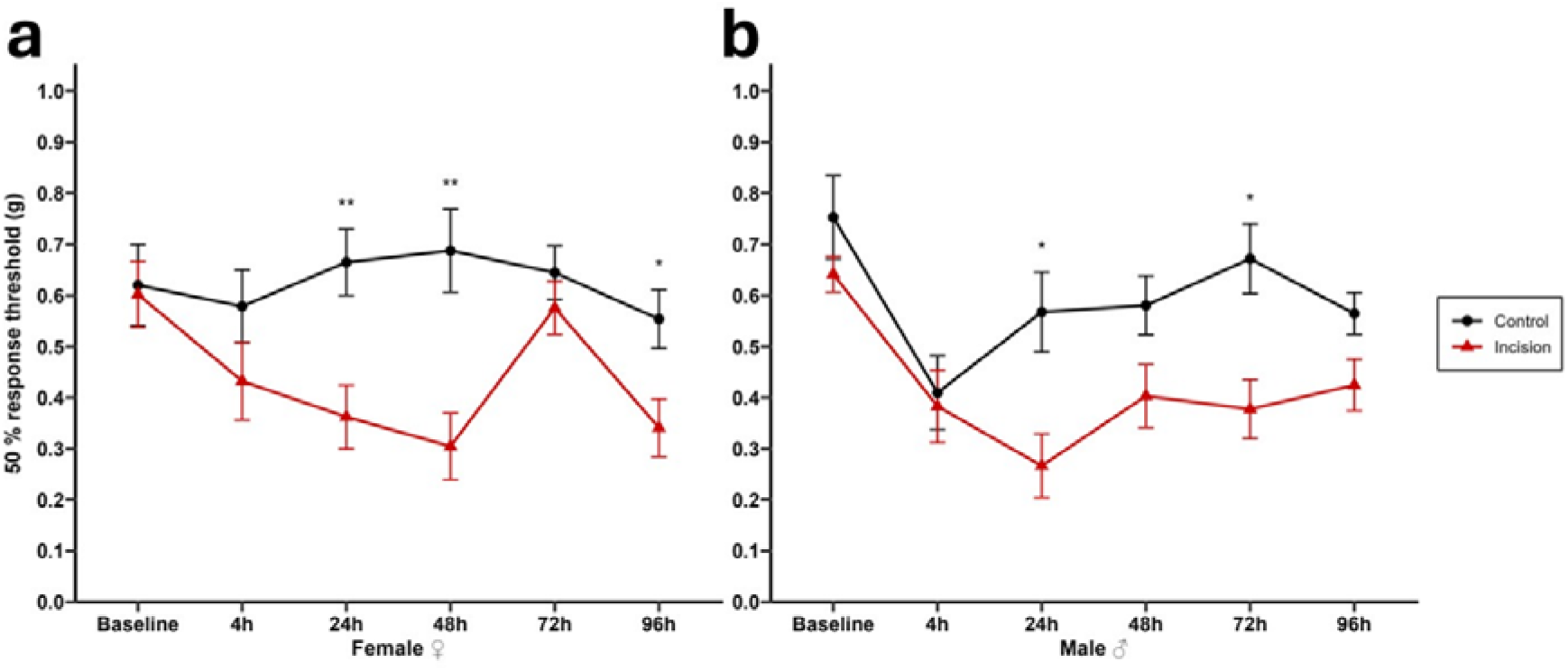
Injury-dependent changes in mechanical sensitivity following plantar incision. **a,** Female mice subjected to plantar incision show reduced 50 % withdrawal thresholds compared with controls in a von Frey assay across the postprocedural time course. **b**, Male mice in both groups initially decreased in withdrawal thresholds at 4 h, whereafter the control mice approached baseline levels, unlike the incision group which showed increased sensitivity. ANOVA revealed a significant main effect of procedure, and stars represent FDR-corrected *post hoc* comparisons. No significant sex- or time-related differences were detected (see results for statistics). Data are shown as mean ± s.e.m., *n* = 12 animals per group. **\* =** P < 0.05, ** = P < 0.01

Two deviations from the overall postprocedural pattern were observed that were not consistent across groups: The female incision group deviated at 72 h from the overall pattern (2a), and a decline was observed at 4 h after anaesthesia in both male groups, including controls (2b).

### Home-cage activity after the procedure

In the postprocedural period, locomotion was significantly reduced in all cages relative to their baseline levels (paired t-test; t(30) = -3.21, P = 0.00315) (Fig. 3a and b). An ANCOVA assessed the effects of procedure (incision/control) and sex (male/female) on postprocedural activity, adjusting for baseline values. Baseline activity was significantly associated with postprocedural activity (F(1, 26) = 17.094, P < 0.001), indicating that cages with higher baseline activity also had higher postprocedural activity. No effect of procedure, sex, nor their interaction was detected. A postprocedural decrease in home-cage activity was also observed in running-wheel activity (Fig. 3c and d) when comparing postprocedural and baseline activity within cages (paired t-test; t(22) = -3.76, P = 0.001); mean running-wheel activity was 6.61 km per day (SD ± 6.15) in the baseline period and fell to 4.41 km (SD ± 4.18) after the procedure. (See Fig. S1 and S3 for detailed activity data). When including incision and sex in an ANCOVA, only the baseline activity was a significant predictor of postprocedural activity (F(1, 18) = 121.662, P < 0.001), while neither sex, incision, nor their interaction were. Given the absence of group differences, we analysed data on a per-day basis to assess temporal patterns following the procedure.

**Fig. 3:**
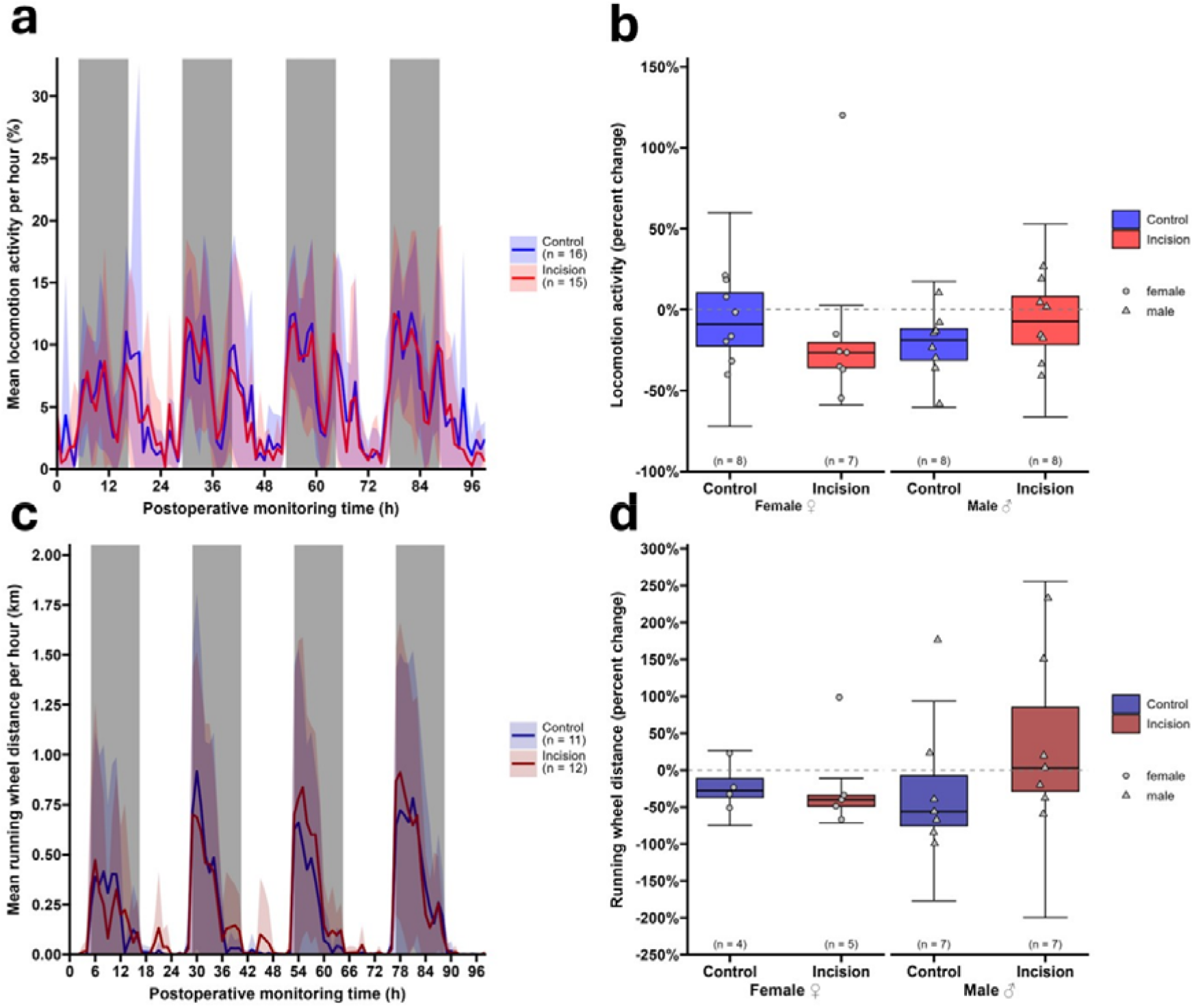
Postprocedural home-cage activity does not differ between groups. **a**, Postprocedural locomotion activity and **c**, running-wheel activity were recorded continuously for 100 hours in the home cage of mice subjected to plantar incision or control procedure. Data are shown as mean ± s.d. (two mice per cage). Grey boxes indicate the facility’s dark phases. **b**, Change in locomotion and **d**, change in running wheel activity were calculated as the sum of the values in **a** and **c**, respectively, relative to the mean daily baseline activity. Boxplots show the median, interquartile range, and whiskers 1.5× the interquartile range in length, with individual data points representing each cage. No differences between procedure groups were detected (see results for statistics). Horizontal dashed lines indicate baseline activity.

### Postprocedural activity per day

We hypothesised that incision-related differences between groups would be most pronounced during the immediate postprocedural period, starting after recovery from anaesthesia and lasting the first full 24 h after the procedure. However, no differences between groups were observed during this period (Fig. S2 and S4), neither for locomotion nor running-wheel activity.

When comparing daily postprocedural activity at the cage-level to baseline (Fig. S1 and S3), independent of incision status, activity was significantly reduced on the first day both for locomotion (t(31) = -3.17, P = 0.0141) and running-wheel activity (t(23) = -5.74, P < 0.001).

Running-wheel activity was still significantly reduced on postprocedural days 2 and 3 (P = 0.013 and P = 0.012, respectively), whereas locomotion activity was not (P = 0.095 and P = 0.11, respectively).

## Discussion

We evaluated whether home-cage monitoring (HCM)-based activity measures can detect pain-related behavioural changes in a plantar incision mouse model of postoperative pain. The model produced increased mechanical sensitivity in the von Frey test (Fig. 2), lasting up to five days after the procedure, consistent with previous work^12^. In our experiment, ‘procedure’ refers to the combined effects of handling, anaesthesia, preparation of surgical site and, where applicable, incision. A transient postprocedural reduction in home-cage locomotion and running-wheel activity at cage level was observed when activity was compared with baseline (Fig. 3). However, spontaneous activity did not differ between incision and control groups, in either sex, under the tested conditions.

The absence of detectable changes in spontaneous activity does not rule out the presence of pain. Rather, it reflects the deliberately mild nature of the paw incision model, which induces postoperative pain-related changes in only one paw. This indicates that cage-level home-cage activity alone is insufficient as an outcome measure when the research question specifically concerns mild or local, pain-related changes. When we used von Frey testing, the incised mice showed significantly increased mechanical sensitivity around the incision site compared to the control group. When using von Frey testing, several inherent limitations must be considered, including experimenter-dependent variability^4^, inconsistent mechanical force of calibrated filaments depending on filament application angle, application speed, bending angle of the filament during application^8^, and, most importantly, the test requires extensive acclimatisation up to several hours prior to each test^13^, which can be stressful for the mice.

Further, in our experiment, von Frey testing could not be performed in a blinded manner, as the experimenter needed to visually identify the plantar surface of the paw, thereby also revealing whether an incision was present or not. This lack of blinding introduces a risk of performance bias, which should be considered when interpreting von Frey thresholds in this model. Taken together, these limitations motivate the use of automated, observer-independent approaches such as home-cage monitoring. Despite these limitations, the assay showed withdrawal thresholds with a clear separation between the groups. Because the overall response pattern matched what is well-documented for the plantar incision model^12, 14, 15^, we therefore consider the von Frey test sufficiently robust for determining the presence of local pain-related changes in this study. Unexpectedly, a decline was observed at 4 h after the procedure in both male groups, including controls (Fig. 2b). This decline is potentially related to stress-induced hyperalgesia from the anaesthetic procedure^7, 16, 17^, emotional contagion between animals ^18^, or simply a reflection of natural biological variation^19^. These speculations cannot be resolved with the present data.

Following general anaesthesia, with or without a plantar incision, home-cage activity declined measurably. We found a transient reduction in home-cage activity of both groups (anaesthesia-only and incision), most pronounced within the first 24 h following the procedure. Locomotion returned to baseline within 24 h after the procedure, whereas running-wheel activity remained reduced for up to 72 h, suggesting greater sensitivity to procedure-related effects. The absence of incision or sex effects indicates that this likely reflects a non-specific, general response to the procedure rather than an incision specific behavioural phenotype under the conditions tested. This demonstrates that the HCM system is capable of detecting broad, non-specific disruptions of normal behaviour induced by the intervention and subsequent recovery. However, this general postprocedural reduction may have masked any additional incision-related differences. While the von Frey test could detect differences in evoked pain-related behaviour, home cage locomotion and running-wheel activity lacked the sufficient sensitivity to detect spontaneous postprocedural pain-related changes. This is despite readouts being unaffected by human interpretation during data acquisition. These findings do not support our hypothesis that HCM-based metrics can detect incision-related changes in this model. Likewise, we found no evidence supporting pain-related changes during the first 24-hour period, therefore our second hypothesis is also rejected. Together, these findings highlight the limitations of cage-level activity metrics as indicators of mild or local postoperative pain.

In higher-intensity pain models, such as prolonged joint-pain^20, 21^, and cancer-induced bone pain^22^, only short-lasting decreases in home-cage activity were found immediately following the model induction, if any were observed at all. Rather, activity patterns changed, with decreased lengths of activity bouts, while overall daily activity rhythms remained stable.

Considering that overall home-cage activity levels appear relatively stable in mild and higher-intensity pain models, cage-level home-cage activity might be too crude to measure pain.

We found, as others have^23, 24^, a decrease in postprocedural HCM-based activity compared with baseline measurements. We could however not find an effect of the experimental incision. No sex-related changes were detected, supporting the hypothesis that incision-related responses would not differ between sexes under these conditions. Previous HCM studies reported no sex differences in naïve C57Bl/6J mice at 9-10 weeks of age. However, sex-related differences emerged after they aged beyond 13-14 weeks, with females displaying higher activity levels than males. Despite the lack of sex differences in naïve mice aged 9-10 weeks, induction of a robust joint-pain model resulted in different responses between the sexes. Specifically, injured female mice maintained their activity levels for longer during the night phase compared with injured males^21^. Our study used mice that were approximately 8-10 weeks old, and the absence of baseline sex differences in activity is consistent with previous findings in young animals. Similarly, the lack of an injury × sex interaction in our study likely reflects the absence of detectable injury effects in this relatively mild pain model.

When evaluating the readouts derived from the HCM-system, we and others^22^ observed that running-wheel activity appears to be the more sensitive measure compared to cage-level locomotion activity. While both measures declined after the procedure, running-wheel activity was reduced for 72 hours, compared to 24 hours for locomotion activity. Presumably, this could be explained by the distance travelled in the running wheel primarily serving a recreational purpose^25^, whereas cage-level locomotion supports fundamental needs like eating, drinking, and nesting. While the experimental procedure altered normal behaviour, recreational behaviour might be among the first to decline during recovery, while other important behaviours are continued despite being under recovery. This finding partially supports our hypothesis, that running-wheel activity is the more sensitive metric under the tested conditions. Yet, the metrics could not be used to discriminate the two procedures, and with no evidence of an incision-related effect, we can neither confirm nor reject our hypothesis.

Several technical and biological factors related specifically to the running-wheel setup may explain the inability to detect incision-related effects in this study. Manzanares et al.^26^ described that upright running wheels may be detrimental to the mice using them, as they require a posture with ventral arching of the spine and hyperflexion of the tail. By contrast, angled running wheels allow a more natural posture that better resembles a mouse’s natural running gait. When mice are given the choice, they prefer an angled wheel, as indicated by increased speed, total running distance, and daily running duration^26^. Activity in an upright running wheel also increases when increasing the wheel size from 13 cm in diameter to 17 cm^27^, suggesting that when less back arching is required, the mice’s willingness to run increases. Unfortunately, we had to use 11 cm vertically oriented wheels to ensure compatibility with the HCM system. If mice are less inclined to use their running wheel, potential pain-related effects will be difficult to detect. Therefore, the wheel design and orientation may have contributed to the low sensitivity of the running-wheel metric in detecting incision-related behavioural changes. In addition, because cages were not handled during the baseline period, bedding and nesting material occasionally obstructed the running wheel. Consequently, baseline running-wheel activity was likely affected in some cages, and eight cages in total were excluded due to low baseline activity. Taken together, these technical and biological constraints must be considered, and they may have contributed to the running-wheels’ inability to detect incision-related effects.

Beyond metric-specific considerations, several broader methodological factors are likely contributors to the limited sensitivity of activity based readouts in this study. These include the mild nature of the pain model, the experimental unit being a cage containing two mice instead of a single individual, and introducing uncontrollable group dynamics that may influence behaviour, such as competition for access to the running wheel. Although co-housing conspecifics may exacerbate painful phenotypes^28^, we did not observe evidence of such an effect in this study. Moreover, overall activity is a crude proxy for animal welfare and may fail to capture subtle, localised, pain-specific behavioural changes. In this context, housing conditions may also play a role: The cellulose bedding used in this study is softer than commonly used hardwood chip or corncob bedding, and harder bedding materials, such as aspen, have been reported to increase sensitivity in evoked assays, including von Frey and Hargreaves tests^6^. It could be argued that harder bedding materials may make injured animals more reluctant to use the affected paw during spontaneous cage activity. Consequently, softer bedding types could reduce the behavioural contrasts between injured and uninjured groups, potentially limiting the sensitivity of home-cage activity measurements. However, the same study reported that running wheel activity was unaffected by bedding type, despite comparing relatively hard aspen bedding with relatively soft TEK-Fresh bedding^6^. This finding suggests that the influence of bedding type may differ between behavioural outcomes, with effects being more readily detected in evoked pain assays than in spontaneous activity measures.

In conclusion, HCM-based metrics must be developed further to reliably detect behavioural changes related to plantar paw incision. Our primary aim was to test whether mild postoperative pain could be detected using non-invasive home-cage readouts. We have shown that the plantar incision model does not produce detectable alterations of spontaneous home-cage behaviour. The running wheel metric proved to be more sensitive than overall cage activity, underlining the need for a continued development of more sophisticated home-cage-derived behavioural measures. These findings emphasise that, while home cage monitoring offers compelling opportunities for continuous, non invasive observation, commonly used activity based metrics alone cannot yet serve as a standalone screening tool for animal welfare or for detecting mild or localised pain related behavioural changes. Future work should focus on integrating higher-resolution behavioural features or individual-level tracking to improve sensitivity to subtle pain-related changes.

## Methods

### Animals and housing

For this study, we used 56 male and 56 female mice. For the von Frey study, 24 male and 24 female specific pathogen free C57BL/6Jrj mice (Janvier; Le Genest-Saint-Isle, France), aged 8 weeks on arrival, were housed in a facility accredited by the Association for Assessment and Accreditation of Laboratory Animal Care (AAALAC) at the Faculty of Health and Medical Sciences, University of Copenhagen. Upon arrival, mice were ear-marked and housed in individually ventilated cage (Blue Line Type III 1290, Tecniplast S.p.A, Buguggiate, Italy), four animals per cage. After seven days of acclimatisation, we transferred the mice to an experimental room for behavioural testing and housed them in open-top cages inside a ventilated cabinet (“Scantainer”, SCANBUR, Karlslunde, Denmark). The mice were housed on soft cellulose bedding (ALPHA-dri, Shepherd Specialty Papers, Watertown, TN, USA), with clear tunnels (50 × 3 × 100 mm Clear Handling Tunnel, Datesand, Bredbury, UK), 2-4 aspen gnawing bricks (Chew Block Small, Tapvei, Harjumaa, Estonia), red transparent plastic shelters (JAKO, Molytex, Glostrup, Denmark) and nesting material (10 g Carfil Nesting Cup, SCANBUR). Mice had *ad libitum* access to water and food pellets (Altromin 1314, Altromin Spezialfutter GmbH & Co. KG, Lage, Germany) throughout the study. These cages contained no running wheels. The facility maintained a temperature of 22 (± 2) with a humidity of 55 % (± 10 %), and the cages were ventilated at 75 air changes per hour. Animal rooms were on a 12/12-hour light/dark cycle with a 30-minute twilight phase, with lights at 50 % intensity from 5:30, increasing to 100 % at 6:00, and decreasing to 50 % at 17:30 before turning off at 18:00. The experimental room was maintained on a 12/12-hour light/dark cycle without the twilight phase.

For the activity study, 32 male and 32 female specific pathogen free C57BL/6Jrj mice (Janvier; Le Genest-Saint-Isle, France), aged 8 weeks on arrival, were housed in the same facility, and were pair housed in individual ventilated cages (EM-500, Tecniplast) equipped with a running wheel (GYM500 Running Wheel, Tecniplast) that had the same environmental conditions as in the von Frey study (bedding, enrichment, temperature, humidity and light cycle, including the twilight light phases). All cages were housed on the same DVC rack, with incision and control cages positioned alternatingly on a single row for each cohort to minimise potential rack-position effects. As changes to bedding can significantly alter home cage behaviour in mice for up to 5 days^29^, we did not change bedding or make major alterations to the home-cage environment during the study.

### Ethics

The Danish Animal Experiments Council approved all procedures (license 2024-15-0201-01627) and the study was overseen by a local animal welfare body. The protocol was not formally preregistered. We defined humane endpoints before initiating the study. They included pronounced dehydration, lethargy, infection of the incision wound, weight loss above 20 %, and blood in urine, mouth or faeces. In addition, we used a scoring system twice daily to monitor acute and accumulated lameness with acute severe lameness and moderate lameness lasting more than three days being humane endpoints. No animals reached the predefined humane endpoints. However, one mouse was euthanised due to surgery related complications before recovering from anaesthesia. Consequently, its cage-mate was euthanised to avoid single housing and incomparable data from this experimental unit.

### Plantar incision model

We anaesthetised all mice with isoflurane. Mice were individually placed in a sliding top chamber (Kent Scientific) with 5 % isoflurane delivered in 100 % oxygen (flow rate of 500 ml/min) until loss of the righting reflex. Anaesthesia was maintained with ∼1.9 % isoflurane delivered in 100 % oxygen via a face mask for spontaneous breathing (flow rate 100 ml/min). The mouse was placed on a heated mat (1180 x 280 mm - 62W, LP Racks, Juelsminde, Denmark) and the right hind paw was cleaned and disinfected *lege artis* with 4 % chlorhexidine soap and 70 % ethanol. No perioperative analgesia was administered, as the study aimed to assess pain-related outcomes, and analgesic intervention would be a confounding factor in the experimental model.

The plantar incision was performed as described by Cowie et al.^14^ with minor changes. After preparation, the paw was incised with a 4 mm longitudinal incision through the skin with a #11 surgical blade. The *flexor digitorum brevis* muscle was lifted with curved forceps and split in half longitudinally. The area was moistened with sterile saline, the muscle was returned to the wound cavity, and the skin was sutured with two interrupted sutures (Ethilon 6-0 nylon monofilament, ref. 697H, Ethicon Inc., Cornelia, GA, USA). Anaesthesia-only mice had their paw cleaned and disinfected under anaesthesia and were kept unconscious for 13 minutes; the average time to complete a surgery. The procedures were conducted between 9:00 and 12:00 in the morning. The procedure was performed once per individual under fully aseptic conditions by an experienced experimenter.

### Von Frey monofilament measurements

To assess mechanical sensitivity, we used von Frey monofilament testing, using the up-down method according to previous methodology^20^, at baseline and 4 h, 24 h, 48 h, 72 h, and 96 h after the procedure (see Fig. 1a). Animals were allocated to groups using matched pairs based on stratified baseline values completely by random without a randomisation sequence. Four mice (two incision and two control animals) were housed per cage.

We placed the mice individually in plastic chambers (96 mm × 96 mm × 140 mm, Enclosure 37000-007, Ugo Basile, Gemonio, Italy) for 60-90 minutes of acclimatisation. Von Frey filaments were carefully applied underneath the right hind paw for five seconds and if no reaction was apparent (flinching, licking paw or withdrawing it, jumping, or toe spreading), we recorded “no response.” A positive response resulted in application of the next lower-force filament. After the first change in response pattern (a “no response”), the next higher-force filament was applied, after which filament force was increased or decreased depending on whether a response occurred, until four applications following the first no response.

Filaments (Semmes-Weinstein set of monofilaments, Ugo Basile) were pre-calibrated with a range from 1.0 gram to 0.008 gram with 0.6 gram as the first stimulus. For calculation of 50 % withdrawal threshold, λ = 0.24 was used in the Chaplan formula: 50 % threshold (g) = 10^(*x*^*^f^*^+ λ^ * *^k^*^)^), where *x_f_* is the log value of the last applied filament, λ is the average log-increment between the applied filaments, and *k* is a value corresponding to the response pattern, see table 7 in ^30^. For this experiment, the experimental unit was each individual mouse (n = 12 mice per sex × procedure group).

### Home-cage monitoring measurements

Two measures were continuously collected from the home-cages at the cage level: Locomotor activity per hour, defined as “locomotion” in percentage, was measured using an electromagnetic plate underneath the cage (see ^31^ for details). Voluntary running-wheel activity was quantified as distance travelled in kilometres per hour. Following acclimatisation (seven days), baseline activity was recorded for five days. For each of the four cohorts, mice were housed n = 2 per cage with a cage mate of the same procedure status (incised with incised, and control with control) with four cages per procedure group per cohort. Procedure allocation was assigned alternatingly, rather than by randomisation. On the day of the procedure, both mice from a cage either received a plantar incision or control procedure, making each cage the experimental unit for these analyses (n = 8 cages per sex × procedure group, except for the female incision group; n = 7 cages due to surgery complications).

Postprocedural activity was recorded for four days (see Fig. 1b and c). Cages were not handled during the acclimatisation and baseline periods. During the postprocedural monitoring period, cages were only handled for welfare monitoring twice daily by briefly removing one cage at a time from the rack.

### Statistics

We analysed data in R (version 4.5.0)^32^ using Rstudio^33^. The complete code and raw data are provided in supplementary information. Statistical significance was defined throughout as P < 0.05. Data were assumed to be normally distributed.

Mechanical sensitivity was analysed using a repeated measures analysis of variance (ANOVA), excluding pre-procedure baseline values, with time as a repeated factor and procedure and sex as between-subject factors. *Post hoc* comparisons at individual time points were performed using two-sided unpaired t-tests, with P values adjusted for multiple comparisons using the Benjamini-Hochberg (BH) procedure^34^ for false discovery rate (FDR) corrections.

Home-cage activity was first normalised to a per-day average for each monitoring period (baseline and postprocedural). For each cage, mean daily activity during the 4-day postprocedural period was compared with mean daily activity during the 5-day baseline period using a two-sided paired t-test on cage-level differences. Effects of procedure and sex were included in an analysis of covariance (ANCOVA), with postprocedural activity as the outcome and baseline activity used as a predictor. The ANCOVA analyses were repeated on a subset of the postprocedural period, covering only the first 24 hours after the procedure. To assess daily postprocedural activity relative to baseline, paired comparisons between baseline and each day following the procedure were performed using two-sided t-tests followed by FDR correction (Benjamini-Hochberg procedure).

One cage was excluded due to surgery-related complications in the home-cage monitoring experiment (female incision group). For running wheel analysis, cages with total activity < 0.1 km during the 5-day baseline period were excluded (n = 5 from control group, n = 3 from incision group).

We did not apply blinding during experimentation or analysis. No *a priori* sample size calculation was performed; sample sizes were constrained by surgical capacity.

## Acknowledgements

Running wheels were kindly provided by the Rodent Metabolic Phenotyping Platform at the Novo Nordisk Foundation Center for Basic Metabolic Research. Thanks to Trine Marie Ahlman Glahder and Daniel Kylmann Hansen for their support during the surgery procedures.

## Contributions

M.M. and K.A. conceptualized the study. M.M., K.A., S.H., and O.K. designed the experiments. M.M. and T.B.B. performed the *in vivo* work. M.M., T.B.B and O.K. processed data. M.M. and O.K. developed code for analysis and figures. M.M. drafted the paper. M.M., T.B.B, P.B., K.A., S.H., and O.K. reviewed and edited the paper and figures. P.B., K.A., S.H., and O.K. supervised the project. P.B. provided funding. All authors have read, reviewed, and agreed to the published version of the paper.

## Supplementary information

Supplementary information is available for this paper. The code used for analyses and generating figures is available here [<u>10.6084/m9.figshare.c.8564660</u>]. Statistical analyses were carried out using R^32^, conducted in RStudio^33^. R-packages used for the statistical analyses are: tidyverse^35^, ggprism^36^, conflicted^37^, and ggh4x^38^. Their respective uses are explained in the code files.

**Fig. S1:**
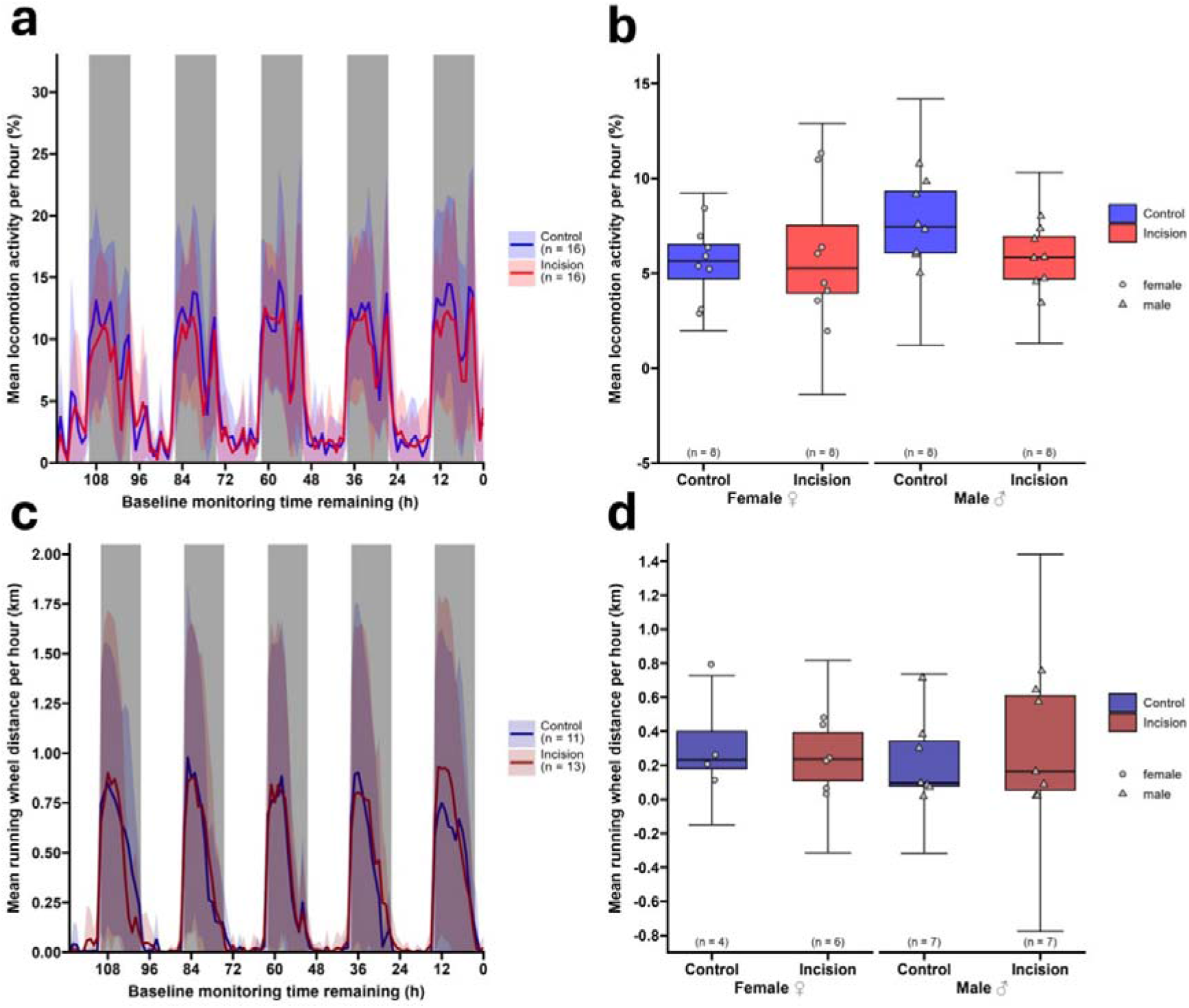
Baseline home-cage activity. **a**, Locomotion activity and **c**, running wheel activity were recorded continuously for 120 hours in the home cage prior to the procedure. Data are shown as mean ± s.d. (two mice per cage). Grey boxes indicate the facility’s dark phases. **b**, Average locomotion activity per hour and **d**, average running-wheel activity per hour were calculated from the corresponding activity in **a** and **c**, respectively. Boxplots show the median, interquartile range, and whiskers 1.5× the interquartile range in length, with individual data points representing each cage. No differences between groups were detected (see results for statistics).

**Fig. S2:**
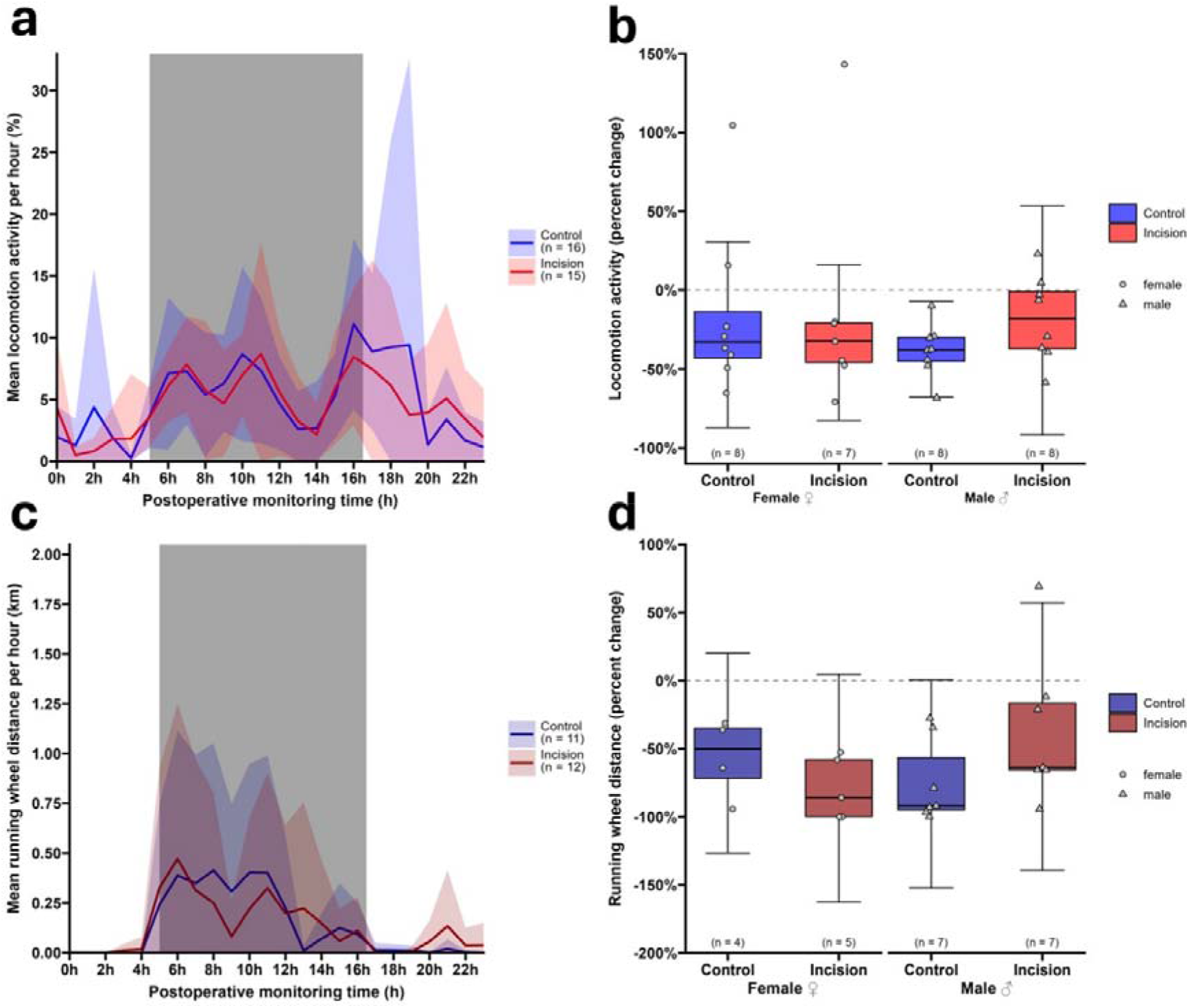
Home-cage activity immediately following the procedure. **a**, Postprocedural locomotion activity and **c**, running-wheel activity during the first 24-hour postprocedural monitoring period, shown as a cropped segment of the 100 h home-cage dataset shown in Fig. 3. Data are shown as mean ± s.d. Grey boxes indicate the facility’s dark phase. Each cage contained two mice. **b**, Percentage change in locomotion activity and **d**, percentage change in running-wheel activity were calculated from the truncated 24-hour period in **a** and **c**, respectively. Values are expressed relative to the mean daily activity in the baseline period. Boxplots show the median, interquartile range, and whiskers 1.5× the interquartile range in length. Individual data points represent each cage. Horizontal dashed lines indicate baseline mean daily activity.

**Fig. S3:**
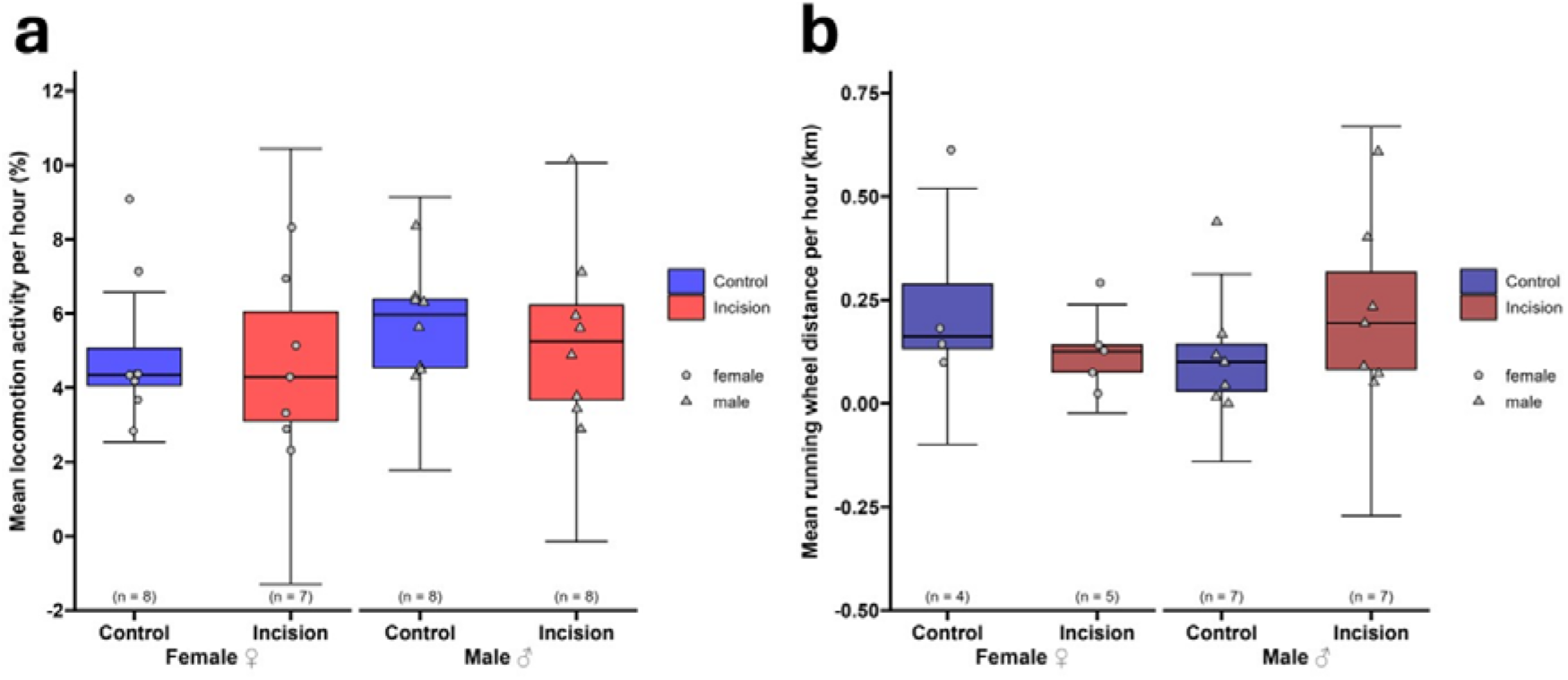
Mean home-cage activity per hour during the entire postprocedural period. **a**, Locomotion activity and **b**, average running-wheel activity per hour of the entire postprocedural period. Activity was recorded continuously in the home cage for mice subjected to plantar incision or control procedure. Data are shown as mean ± s.d. (two mice per cage).

**Fig. S4:**
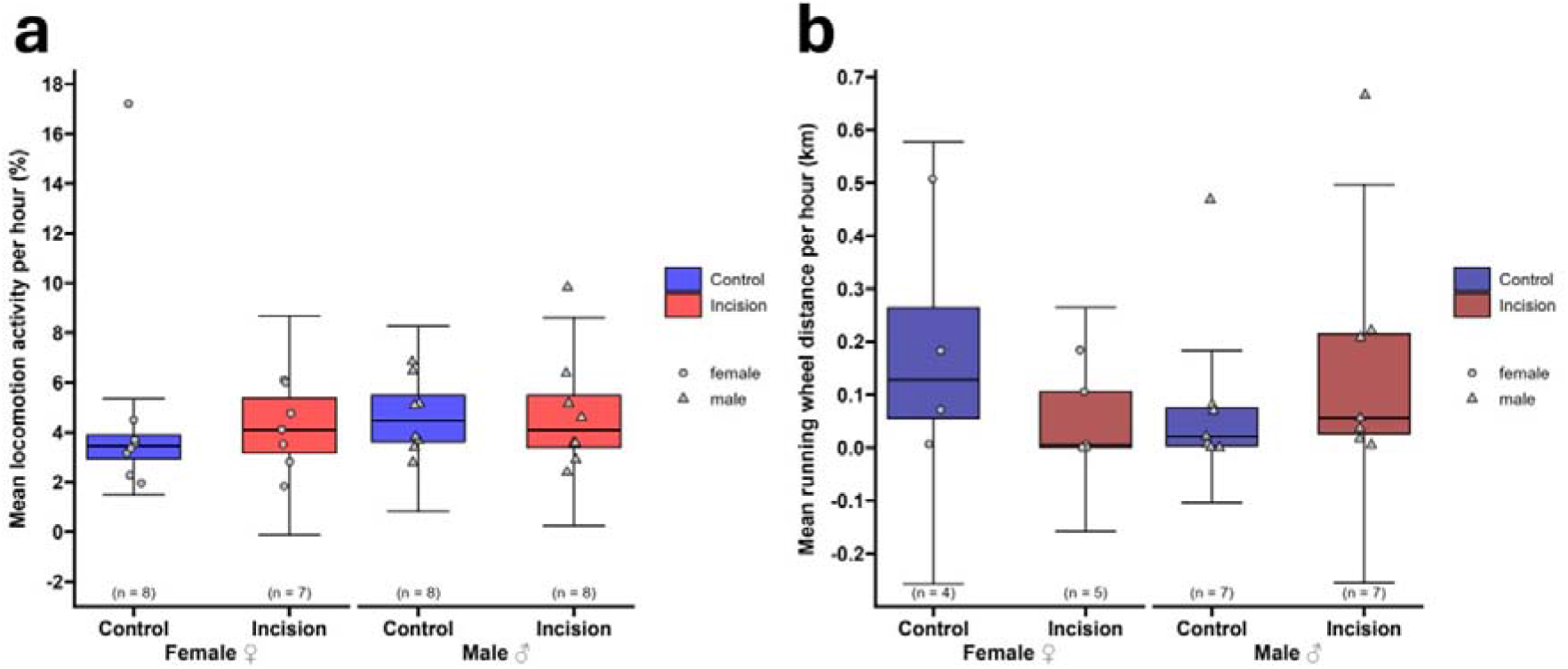
Mean home-cage activity per hour during the 24-hour postprocedural period. **a**, Locomotion activity and **b**, running-wheel activity per hour during the 24-hour postprocedural period. Activity was recorded continuously in the home cage for mice subjected to plantar incision or control procedure. Data are shown as mean ± s.d. (two mice per cage).

